# First report of *Telenomus remus* parasitizing *Spodoptera frugiperda* and its natural parasitism in South China

**DOI:** 10.1101/697920

**Authors:** Yong-Lin Liao, Bin Yang, Miao-Feng Xu, Wei Lin, De-Sen Wang, Ke-wei Chen, Hua-Yan Chen

**Affiliations:** Institute of Plant Protection, Guangdong Academy of Agricultural Science, Guangdong Provincial Key Laboratory High Technology for Plant Protection, Guangzhou, 510640, China; State Key Laboratory for Biology of Plant Diseases and Insect Pests, Institute of Plant Protection, Chinese Academy of Agricultural Sciences, Beijing 100193, China; Technical Center of Gongbei Customs District, Zhuhai, 519001, China; College of Agriculture, South China Agricultural University, Guangzhou, 510642, China; State Key Laboratory of Biocontrol, School of Life Sciences, Sun Yat-sen University, Guangzhou, 510275, China

**Keywords:** *Telenomus remus*, *Spodoptera frugiperda*, egg parasitoid, biological control

## Abstract

The fall armyworm, *Spodoptera frugiperda*, is a lepidopteran pest that feeds on many economically important cereal crops such as corn, rice, sorghum, and sugarcane. Native to the Americas, it has become a serious invasive pest in Africa and Asia. Recently, this pest was found invaded China and has spread exceptionally fast across the country. As *S. frugiperda* will most likely become a major pest in China, Integrated Pest Management strategies, including biological control methods, should be developed to win the battle against it. Here, we report the detection of *Telenomus remus* parasitizing *S. frugiperda* eggs in cornfields in South China based on morphological and molecular evidence. Our preliminary surveys indicated that the natural parasitism rates of *T. remus* on *S. frugiperda* could reach 30% and 50% for egg masses and per egg mass, respectively. Further application of *T. remus* against *S. frugiperda* in biological control programs are discussed.

## Introduction

The fall armyworm, *Spodoptera frugiperda* (Smith) (Lepidoptera: Noctuidae), which originating from the tropical and sub-tropical of North, Central, and South America, has become a invasive pest of cereals in Africa, India, Myanmar, Thailand, etc, where it has caused serious damage (Kenis *et al*., 2019). This species was first detected in the southeast province of China, Yunnan, in January, 2019, has quickly spread from the south to the north and has now found in 18 provinces (Jiang *et al*., 2019; Cui *et al*., 2019). Potential geographic distribution models have indicated that large parts of China are suitable for the survival of this devastating pest (Lin *et al*., 2019).

In its native and introduced ranges *S. frugiperda* feeds on a wide range of crops. Over 350 different host plants in numerous families have been recorded, with almost 40% of them are economically important (Montezano *et al*., 2018). It is estimated that, just for corn, rice, sorghum and sugarcane, this pest could cause up to $US13 billion per annum in crop losses in Africa (Day *et al*., 2017). Given that corn, sugarcane, and rice are widely grew in South China, *S. frugiperda* will most likely establish as a major pest in this regions (Wang *et al*., 2019). Currently, chemical control is still the main strategy against this pest in China, although some biological control experiments using predators (*Picromerus lewisi* Scott) have been conducted in the lab (Tang *et al*., 2019). However, in the long run, more biological control strategies should be adopted to against *S. frugiperda* under the perspectives of Integrated Pest Management (IPM).

Among over 150 parasitoid species that attack *S. frugiperda*, the egg parasitoid species *Telenomus remus* Nixon (Hymenoptera: Platygastridae) seems to be a promising biological control candidate (Kenis *et al*., 2019). *T. remus* was reported to attack eggs of various *Spodoptera* species in China (Chou, 1987; Tang *et al*., 2010). We considered that this species may be eventually found attack *S. frugiperda* in China as well. In this study, we report the detection of *T. remus* parasitizing *S. frugiperda* eggs and its parasitism in cornfields in South China.

## Materials and methods

### Field sampling

Eggs of *S. frugiperda* were collected from cornfields of three sites in Guangzhou and Foshan, China, in May and June, 2019 (Table 1). Egg masses were brought back to laboratory and individual egg mass was placed in a 10 cm glass tube and kept in a growth chamber set at 26°C±1°C, 40–60% humidity and a 12L:12D light cycle, and checked daily for emergence of *S. frugiperda* or parasitoids. Larvae of *S. frugiperda* from the non-parasitized eggs in the same egg mass usually emerged first and were transferred to artificial diet (originally designed for rearing the tobacco cutworm, *Spodoptera litura* (Fabricius)) and reared until adult to confirm the identification of the hosts and for other uses. Any parasitoids that emerged were placed in 100% ethanol for both morphological and molecular analyses. Eggs of each egg mass, emerged larvae, parasitism and sex ratio of the parasitoids were counted.

**Table 1.** Details of the sampling, numbers, and accession numbers of parasitoids sequenced

| Locality (City) | Coordinates | Collection Date | Host Plant | No. Egg Mass | No. Barcoded | Code & Accession Number |
| --- | --- | --- | --- | --- | --- | --- |
| Huadu District, Guangzhou | 23°29'6.11"N<br>113°16'11.99"E | 8.vi.2019 | corn | 28 | 2 | Huadu_F (MN123241)<br>Huadu_M (MN123242) |
| Panyu District, Guangzhou | 23°3'21"N<br>113°24'41"E | 7.vi.2019 | corn | 5 | 2 | Panyu_F (MN123243)<br>Panyu_M (MN123244) |
| Gaoming, Foshan | 22°48'22.28"N<br>112°34'19.83"E | 18.vi.2019 | corn | 3 | 2 | Gaoming_F (MN123239)<br>Gaoming_M (MN123240) |

### Species identification

Species of *Telenomus* were determined using the characters of Johnson (1984). Considering the similarity of females between *Telenomus* species, genitalia of males collected from each locality was examined. To confirm morphological identifications, genomic DNA was extracted from individual female and male collected from different locality using a nondestructive DNA extraction protocol as described in Taekul *et al*. (2014). Voucher specimens for all molecular data are deposited in the Museum of Biology at Sun Yat-sen University, Guangzhou, China. Following extraction the “barcode” region of the mitochondrial cytochrome oxidase subunit 1 (*COI*) was amplified using the LCO1490/HCO2198 primer pair (Folmer *et al*., 1994). Polymerase chain reactions (PCRs) were performed using Tks Gflex™ DNA Polymerase (Takara) and conducted in a T100™ Thermal Cycler (Bio-Rad). Thermocycling conditions were: an initial denaturing step at 94°C for 1 min, followed by 5 cycles of 98°C for 10s, 45°C for 15s, 68°C for 30s; 35 cycles of 98°C for 10s, 52°C for 15s, 68°C for 30s and an additional extension at 68°C for 5 min. Amplicons were directly sequenced in both directions with forward and reverse primers on an Applied Biosysttems (ABI) 3730XL by Sangon Biotech (Shanghai, China). Chromatograms were assembled with Sequencing Analysis 6 (ThermoFisher Scientific, Gloucester, UK). All the amplified sequences were deposited into GenBank (accession numbers see Table 1).

Sequences obtained in this study were compared with those analyzed by Kenis *et al*. (2019). Sequences were aligned by codons using MUSCLE implemented in MEGA6 (Tamura *et al*., 2013). The alignment was then analyzed using RAxML as implemented in Geneious 11.0.3 with *Gryon cultratum* Masner and *Gryon largi* (Ashmead) (Hymenoptera: Platygastridae) used as outgroups to root the tree.

### Photography

Images of live specimens were captured using a Keyence VHX-6000 digital microscope. Images of mounted specimens were produced with Combine ZP and AutoMontage extended-focus software, using a JVC KY-F75U digital camera, Leica Z16 APOA microscope, and 1X objective lens.

## Results

Only one parasitoid species, *T.* remus, emerged from the 36 egg masses collected from the three sites. The specimens of *T. remus* (Figure 1) collected from the three sites show no morphological variation and match well with the description of this species developed by Nixon (1937) and Chou (1987). The *COI* sequences were identical among females and males sampled from the three collecting sites, and over 99% of the pairs of bases were identical to a series of sequences labeled as *T. remus* available from the Barcoding of Life Data system and the GenBank database. Phylogenetic analysis based on *COI* sequences generated from this study and those used by Kenis *et al*. (2019) showed that the six specimens collected from the three sites of South China were grouped well within the clade of *T. remus* specimens collected from Asia, Africa, and the Americas (Figure 2).

**Figure 1.**
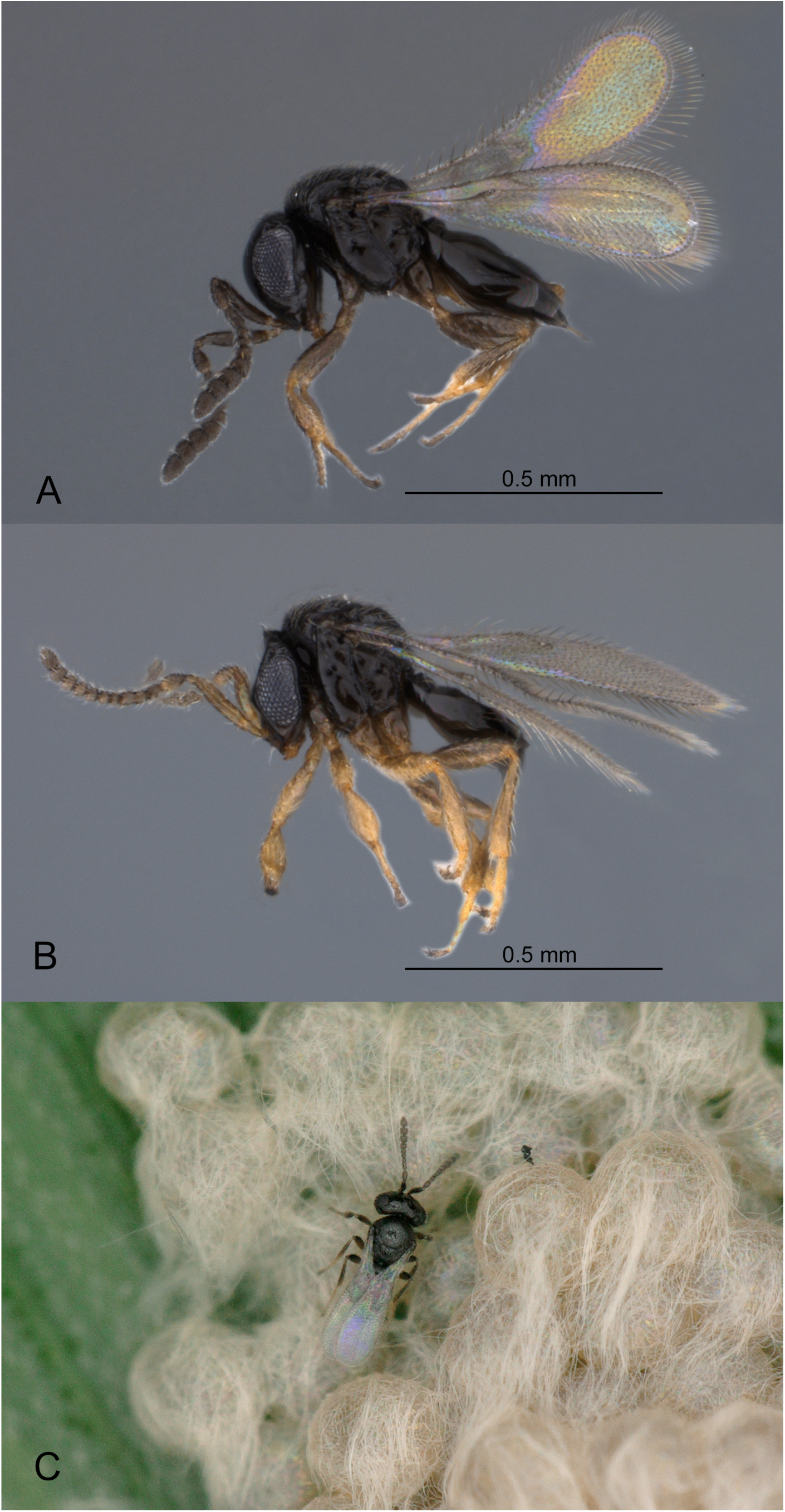
*Telenomus remus* Nixon. **A** Female, lateral habitus **B** Male, lateral habitus **C** A female on egg mass of *Spodoptera frugiperda*.

**Figure 2.**
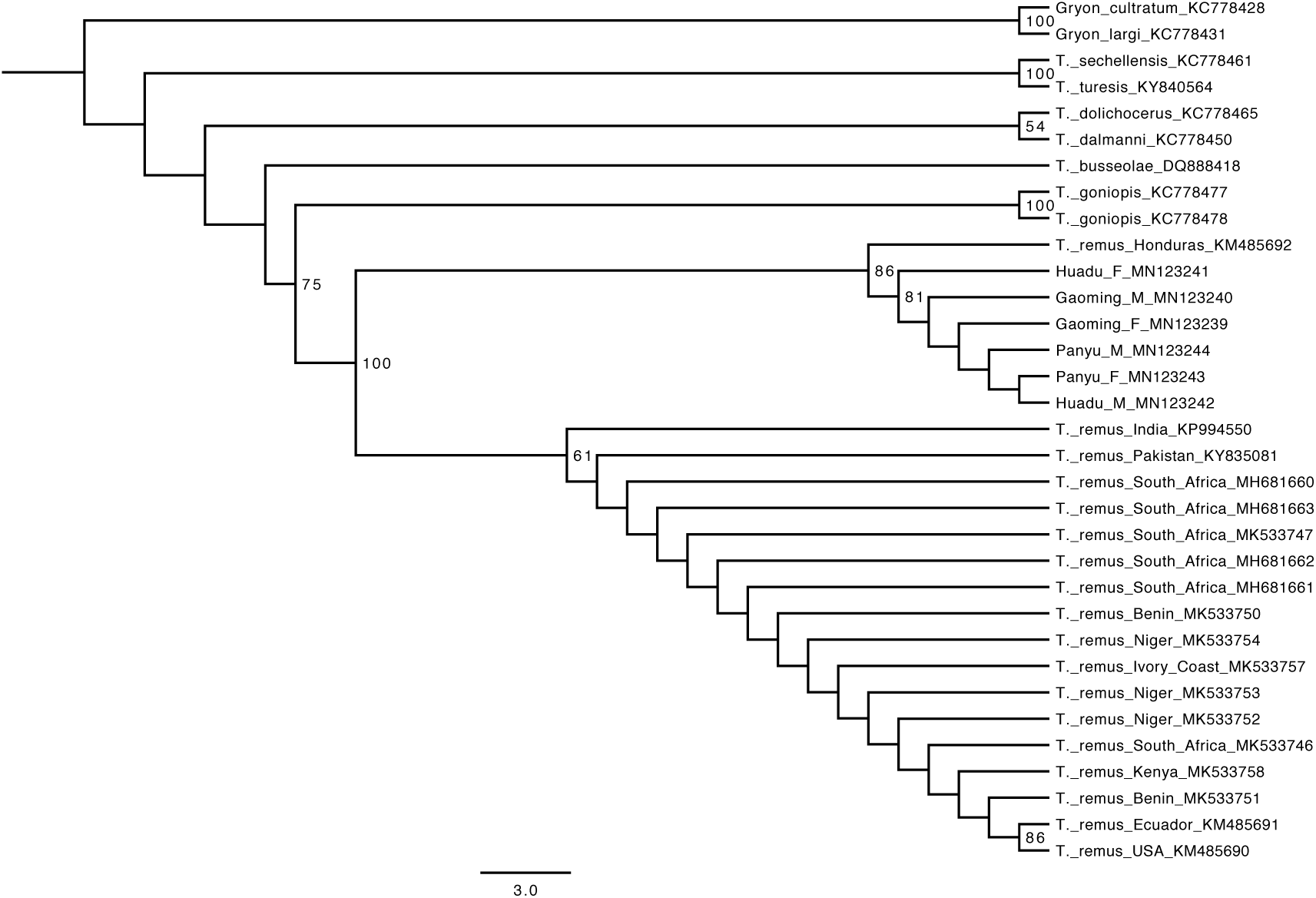
Phylogenetic analysis of *Telenomus remus* and related species by maximum likelihood method based on *COI* sequences. The six sequences generated from this study are indicated with codes and GenBank accession numbers (see Table 1). Boostrap values above 50 indicated on branches.

Twenty-eight, five, and three egg masses of *S. frugiperda* were collected from Huadu, Panyu, and Gaoming, respectively (Table 1). Of the 36 egg masses collected, 11 egg masses (30.6%) were parasitized by *T. remus*. For the 28 egg masses collected from Huadu, we counted the number of layers of each egg mass, parasitism per egg mass and parasitoid sex ratio in details (Figure 3 and Table 2). Of the 28 eggs masses, one-layer and two-layer egg masses seems to be dominant (both 13), followed by three-layer egg masses (3). Of the 7 parasitized egg masses, 6 egg masses were two-layer and one was one-layer. The number of eggs of each parasitized egg masses ranges from 64 to 163. The number of emerged *T. remus* adult from these parasitized egg masses ranges from 29 to 87, with an approximate 79.2%±2.14 female ratio, resulting an approximate 50.86%±2.24 parasitism per egg mass.

**Table 2.**
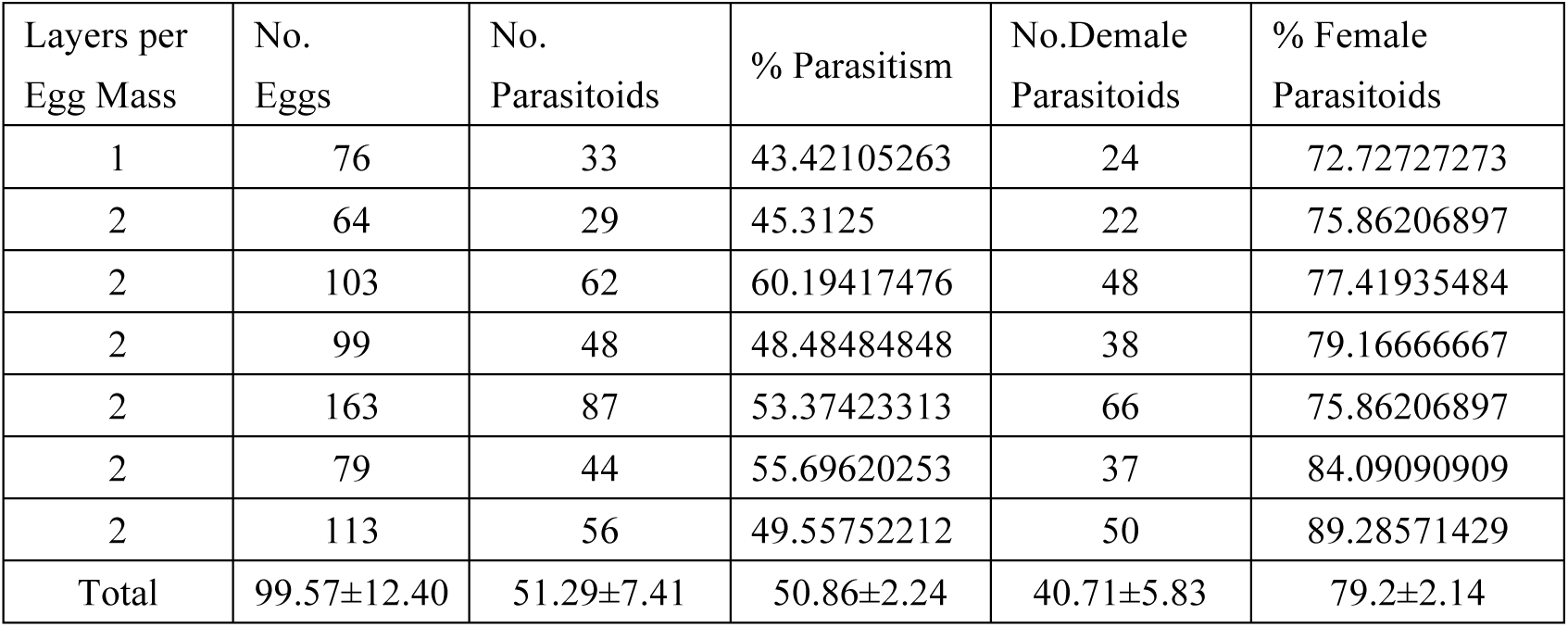
Status of parasitized *Spodoptera frugiperda* egg masses, natural parasitism and sex ratio of *Telenomus remus* collected from Huadu, Guangzhou

| Layers per Egg Mass | No. Eggs | No. Parasitoids | % Parasitism | No. Demale Parasitoids | % Female Parasitoids |
| --- | --- | --- | --- | --- | --- |
| 1 | 76 | 33 | 43.42105263 | 24 | 72.72727273 |
| 2 | 64 | 29 | 45.3125 | 22 | 75.86206897 |
| 2 | 103 | 62 | 60.19417476 | 48 | 77.41935484 |
| 2 | 99 | 48 | 48.48484848 | 38 | 79.16666667 |
| 2 | 163 | 87 | 53.37423313 | 66 | 75.86206897 |
| 2 | 79 | 44 | 55.69620253 | 37 | 84.09090909 |
| 2 | 113 | 56 | 49.55752212 | 50 | 89.28571429 |
| Total | 99.57±12.40 | 51.29±7.41 | 50.86±2.24 | 40.71±5.83 | 79.2±2.14 |

**Figure 3.**
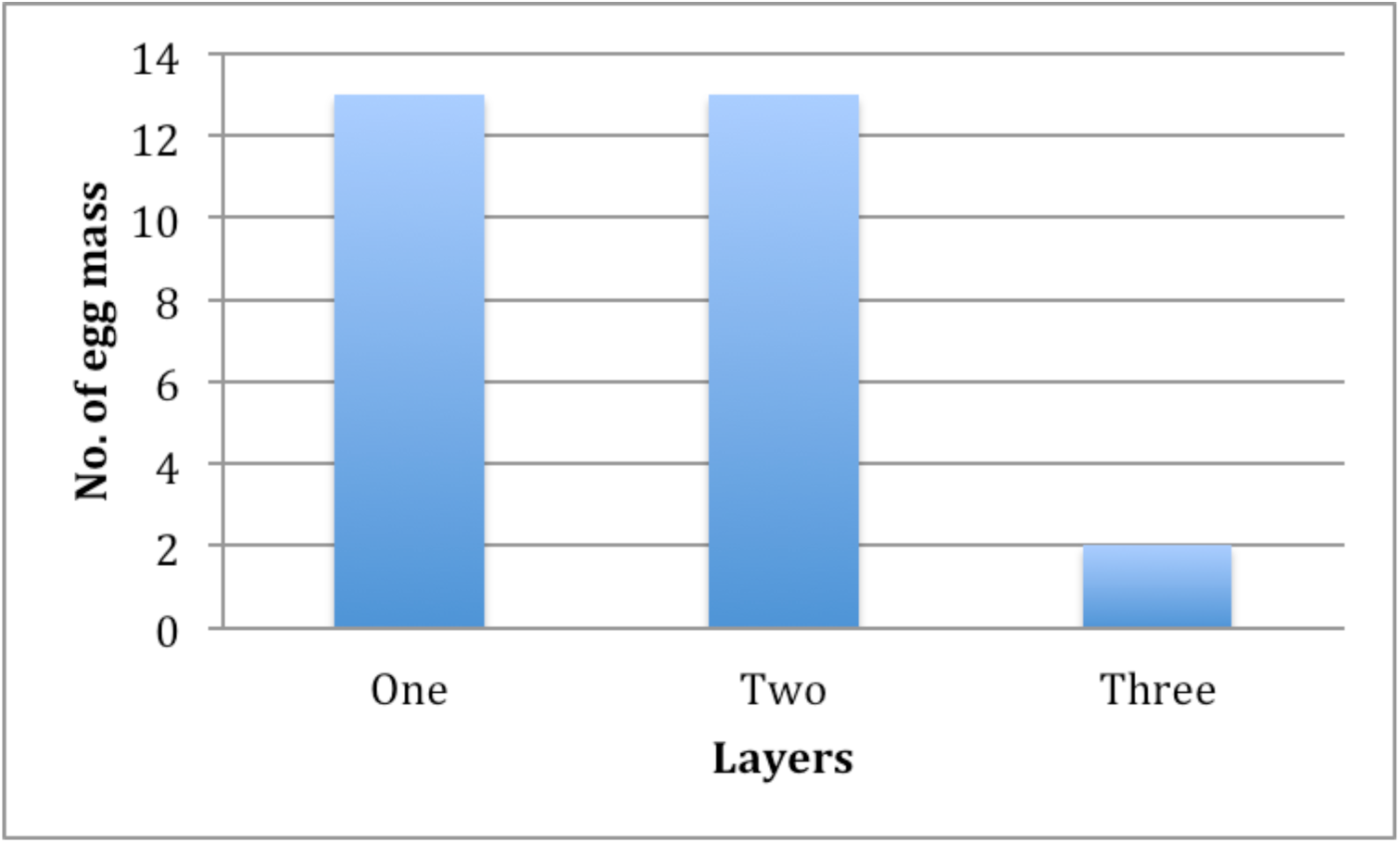
Frequnces of layers of *Spodoptera frugiperda* egg mass.

## Discussion

Both morphological and molecular analyses in this study showed that a *Telenomus* species we found attacking *S. frugiperda* eggs in South China is the promising biological control agent, *T. remus*. In China, this parasitoid species was once reported attack eggs of *Spodoptera* spp. such as *S. litura* and *Spodoptera exigua* (Hübner) (Chou 1987; Tang et al. 2010), but its identity had never been well confirmed. Both of our morphological and molecular data confirmed the presence of *T. remus* and now its parasitism on *S. frugiperda* eggs in China. *Telenomus* spp. are small and morphologically simplified, rendering them difficult to distinguish and identify. However, in the case of using *T. remus* against *Spodoptera* pests, biological control practitioners should note that the colors of legs are just variations between female and male (confirmed by *COI* sequences from both sexes), rather than different species. We recommend that an integrated taxonomy strategy should be applied to identify *Telenomus* species, especially in the case of introducing a parasitoid species into a new region for biological control of pests.

Eggs of *S. frugiperda* are usually laid in masses of approximately 100-200 eggs which are laid in one to multiple layers on the surface of the leaf (Guo et al., 2019). And the egg mass is usually covered with a felt-like layer of scales (setae) from the female abdomen. Our preliminary observation showed that *S. frugiperda* females usually lay one-layer and two-layer egg masses and rarely three-layer in cornfields (Figure 3). Although our current data do not allow us to analyze the preference of different egg masses attacked by *T. remus* due to small sample size (Table 2), this parasitoid species seems to be able to parasitize multiple-layer egg masses. But whether architecture of *S. frugiperda* egg masses affects parasitism rates of *T. remus* requires further investigation. This information is crucial when *S. frugiperda* eggs are used for mass rearing of *T. remus* both for experiments in laboratory or release in the field.

Studies have shown that *T. remus* has great potential use in augmentative biological control against *S. frugiperda* in the field (Cave, 2000). Our observations showed that natural parasitism rates of *T. remus* in corn fields in South China could be up to 30% and 50% for egg masses and per egg mass, respectively. A study conducted in cornfields in Venezuela, parasitism rates of *T. remus* on *S. frugiperda* reached 90% through inundative release in corn cultivation areas (Ferrer, 2001). Further research programs, such as long-term monitoring of natural parasitism and mass release in the filed, should be developed for *T. remus* to evaluate its impact on *S. frugiperda* populations in China.

## Acknowledgments

This study was supported by grants from the Optimization and Popularization of Plant Central Hospital’s Information Platform for Remote Diagnosis and Prevention of Diseases and Pests (2016B080802004) and China Postdoctoral Science Foundation (2019M653187).

## References

Cave, R.D. (2000) Biology, ecology and use in pest management of Telenomus remus. Biocontrol News and Information, 21, 21–26.

Chou, L. Y. (1987) [Note on Telenomus remus (Hymenoptera: Scelionidae).] Bulletin of Society of Entomology, National Chung-Hsing University, 20, 15–20.

Cui, L., Rui, C. H., Li, Y. P., Wang, Q. Q., Yang, D. B., Yan, X. J. et al. (2019) Research and application of chemical control technology against *Spodoptera frugiperda* (Lepidoptera: Noctuidae) in foreign countries. Plant Protection, DOI: 10.16688/j.zwbh.2019300

Day, R., Abrahams, P., Bateman, M., Beale, T., Clottey, V., Cock, M. et al. (2017) Fall armyworm: Impacts and implications for Africa. Outlooks on Pest Management, 28, 196–201.

Ferrer, F. (2001) Biological of agricultural insect pest in Venezuela; advances, achievements, and future perspectives. Biocontrol News and Information, 22, 67–74.

Folmer, O., Black, M., Hoeh, W., Lutz, R., and Vrijenhoek, R. (1994) DNA primers for amplification of mitochondrial cytochrome c oxidase subunit I from diverse metazoan invertebrates. Molecular Marine Biology and Biotechnology, 3, 294–299.

Guo, J. F., Jing, D. P., Tai, H. K., Zhang, A. H., He, K. L., and Wang, Z. Y. (2019) Morphological characteristics of *Spodoptera frugiperda* in comparison with three other lepidopteran species with similar injury characteristics and morphology in cornfileds. Plant Protection, 45, 7–12.

Jiang, Y. Y., Liu, J., and Zhu, X. M. (2019) Occurrence and trend of *Spodoptera frugiperda* invasion in China. Plant Protection, 39, 33–35.

Johnson, N. F. (1984) Systematics of Nearctic *Telenomus*: classification and revisions of the *podisi* and *phymatae* species groups (Hymenoptera: Scelionidae). Bulletin of the Ohio Biological Survey, 6, 1–113.

Kenis, M., du Plessis, H., Van den Berg, J., Ba, M. N., Goergen, G., Kwadjo, K. E. et al. (2019) *Telenomus remus*, a candidate parasitoid for the biological control of *Spodoptera frugiperda* in Africa, is already present on the continent. Insects, 10, 92.

Lin, W., Xu, M. F., Quan, Y. B., Liao, L., and Gao, L. (2019) Potential geographic distribution of *Spodoptera frugiperda* in China based on MaxEnt model. Plant Quarantine. http://kns.cnki.net/kcms/detail/11.1990.s.20190422.1026.002.html.

Montezano, D. G., Sosa-Gómez, D. R., Roque-Specht, V. F., Sousa-Silva, J. C., Paula-Moraes, S. V., Peterson, J. A. et al. (2018) Host plants of *Spodoptera frugiperda* (Lepidoptera: Noctuidae) in the Americas. African Entomology, 26, 286–300.

Nixon, G. E. J. (1937) Some Asiatic Telenominae (Hym., Proctotrupoidea). Annals and Magazine of Natural History, 10, 444–475.

Taekul, C., Valerio, A. A., Austin, A. D., Klompen, H., and Johnson, N. F. (2014) Molecular phylogeny of telenomine egg parasitoids (Hymenoptera: Platygastridae s.l.: Telenominae): evolution of host shifts and implications for classification. Systematic Entomology, 39, 24–35.

Tamura, K., Stecher, G., Peterson, D., Filipski, A., and Kumar, S. (2013) MEGA6: Molecular Evolutionary Genetics Analysis version 6.0. Molecular Biology and Evolution, 30, 2725–2729.

Tang, Y. L,. Chen, K. W., and Xu, Z. F. (2012) Study on Ontogenesis of *Telenomus remus* Nixon (Hymenoptera: Scelionidae). Journal of Changjiang Vegetables, 18, 1–3.

Tang, Y. T., Wang, M. Q., Chen, H. Y., Wang, Y., Zhang, H. M., Chen, F. S. et al. (2019) Predation and behavior of *Picromerus lewisi* Scott to *Spodoptera frugiperda* (J. E. Smith). Chinese Journal of Biological Control, DOI: 10.16409/j.cnki.2095-039x.2019.04.005

Wang, L., Chen, K. W., Zhong, G. H., Xian, J. D., He, X. F., Lu, and Y. Y. (2019) Progress for occurrence and management and the strategy of the fall armyworm *Spodoptera frugiperda* (Smith). Journal of Environmental Entomology. http://kns.cnki.net/kcms/detail/44.1640.Q.20190523.1748.004.html

